# Exploring the Spatiotemporal Influence of Climate on American Avian Migration with Random Forests

**DOI:** 10.1101/2023.03.27.534441

**Authors:** I. Avery Bick, Vegar Bakkestuen, Marius Pedersen, Kiran Raja, Sarab Sethi

## Abstract

Birds have adapted to climatic and ecological cycles to inform their Spring and Fall migration timings, but anthropogenic global warming has affected these long-establish cycles. Understanding these dynamics is critical for conservation during a changing climate. Here, we employ a modeling approach to explore how climate spatiotemporally affects bird occurrence on eBird surveys. Specifically, we train an ensemble of multivariate and multi-response random forest models on North and South American climate data, then predict eBird survey occurrence rates for 41 migrating passerine bird species in a Northeastern American ecoregion from 2008-2018. In October, when many passerines have begun their southward winter migration, we achieve more accurate predictions of bird occurrence using lagged climate features alone to predict occurrence. These results suggest that analyses of machine learning model metrics may be useful for identifying spatiotemporal climatic cues that affect migratory behavior. Lastly, we explore the application and limitations of random forests for prediction of future bird occurrence using 2021-2040 climate projections.

## Introduction

Across the globe, birds undertake Spring migrations to breeding grounds and Fall migrations to winter in warmer climates. In the Western Hemisphere specifically, many bird species migrate northward from their winter homes in the Southeastern USA, the Caribbean, Mexico, and Central America to breeding grounds in the Northern forests of Canada and the United States each Spring, then again southward in the Fall ^1,2^. Bird species rely on climatic and ecological conditions such as temperature, pressure, precipitation, and food availability to trigger the timings of their migrations ^3–6^ in addition to their endogenous rhythms ^7–9^. However, these climatic and ecological triggers are shifting with climate change. In North America, these are projected to be characterized by warming of northern breeding regions, increased rainfall during Spring migration, and less rainfall in southern regions during non-breeding season ^10^.

The broad effects of climate change on migrations have already been demonstrated: shifts to earlier Spring first arrival dates ^2,6,7,11,12^, smaller and more dispersed migratory cohorts ^8^, and a northward shifts in breeding grounds^13^. Previous modeling work based on citizen science data has demonstrated that the climatic suitability of breeding and wintering grounds may be shifting for migrating bird species in South America, with significant decreases in suitable habitat for long-distance migrants in particular ^14^. Indeed, bird species will have to adapt behavioral patterns more quickly than has been done before anthropogenic climate change began^2^.

However, the sensitivities to environmental and ecological cues have been genetically shaped by natural selection to correctly time behaviors such as egg laying to the availability of food, and species often over- or under-correct their timings ^5,15^. By understanding the climatic factors underlying changing bird migration behavior, we can inform species distribution models and conservation practices in a changing climate.

Many studies have explored relationships between migration timings and climatic features through linear-based models, finding that bird migratory rate and arrival time are correlated with climate. For example, warmer Spring temperatures have been linked to earlier migratory arrivals at breeding grounds ^7,8^, and climate in migration stopovers has been shown to affect autumn migration timing^16,17^. These papers have identified ecological trends from post-hoc linear analyses^7,8,17^, but fewer papers use a multivariate modeling approach to directly predict bird species occurrence from climate features^18^. This approach benefits from the ability to use machine learning models, which can leverage non-linear and complex trends in ecoclimatic data ^18–20^ and give better predictions than typical regression models^19^.

Random forests^21^ are a type of decision tree machine learning model that have been successfully applied in ecological settings for regression or classification tasks to relate ecoclimatic features to biodiversity ^20,22^, animal movement and migration ^18^, and species distribution modeling ^23–25^. Random forests use bootstrap sampling to randomly subset training data many times (with replacement). For each bootstrap sample, a decision tree is created by recursively splitting input features to maximize homogeneity of the response features within the output nodes–this is typically determined by mean squared errors within each node for regression problems^26^. This process creates an ensemble of decision trees, each trained on different subsets of feature data, whose predictions are then averaged together. Random forests can be extended to multi-response settings, where splits are optimized by the homogeneity of output nodes across all target variables^27^.

Through averaging the predictions of many weak learners, random forests are well-equipped to handle non-linearity^19^, many interacting features^18^, and prevention of overfitting.^21^ These capabilities are particularly relevant in ecological settings, where many ecoclimatic variables correlate and can influence the presence or behavior of one or more species in non-linear ways^20^. Random forests also have built-in feature importance metrics, which have been applied in understanding which ecoclimatic features best inform predictions of bird species richness^20^. Previous work has demonstrated that purpose-built neural networks^28^ and hierarchal models^29^ can apply dozens of spatial ecoclimatic features to accurately predict occurrence of multiple species. Still, random forests are well-established, high performing, simple to implement, and have built-in feature important metrics, making them an ideal candidate for demonstrating and explaining the capabilities of machine learning models in ecological settings.

In this paper, we apply random forests to predict the occurrence of bird species in a forested North American ecoregion using intercontinental climate data. We first explore whether random forests can successfully integrate spatial climate data to predict many species simultaneously. We next run random forests with lagged climate data, noting model performance and feature importance metrics. In this way, we explore whether climate data in previous months leads to better prediction of current species occurrence and whether the models identify the local climate as more important for prediction than the climate in more distant areas.

Specifically, we use a multivariate and multi-response random forest (MRF) ensemble to predict eBird ^30^ checklist occurrence rates of common migrating passerine species in forests within a Northeastern American ecoregion^31^. Each model incorporates up to 45 spatial climate features across North and South America to predict mean monthly checklist occurrence rates for 41 species across our ecoregion. Lastly, we train a model on historical data, then project future checklist occurrences using several climate models projections averaged from 2021-2040, comparing our results to the general literature consensus of climate impacts on migration timing and discussing the limitations of random forest models for prediction of future trends.

## Methods

### Study Area

We focus our analyses within the Level 2 Ecoregion “Atlantic Highlands” within the Level 1 “Northern Forests” Ecoregion (Fig 1) ^32^. These Environmental Protection Agency (EPA) Ecoregions seek to aggregate areas of similar plant and animal species, as well as abiotic characteristics such as water availability. The “Atlantic Highlands” subregion is characterized by warm summers and cold winters, with higher elevations, more rugged terrain, and less human population than neighboring regions ^31^. Forest type transitions from boreal in the north to deciduous in the southern areas ^31^. Additionally, the region has extensive coverage by eBird surveys.

**Figure 1:**
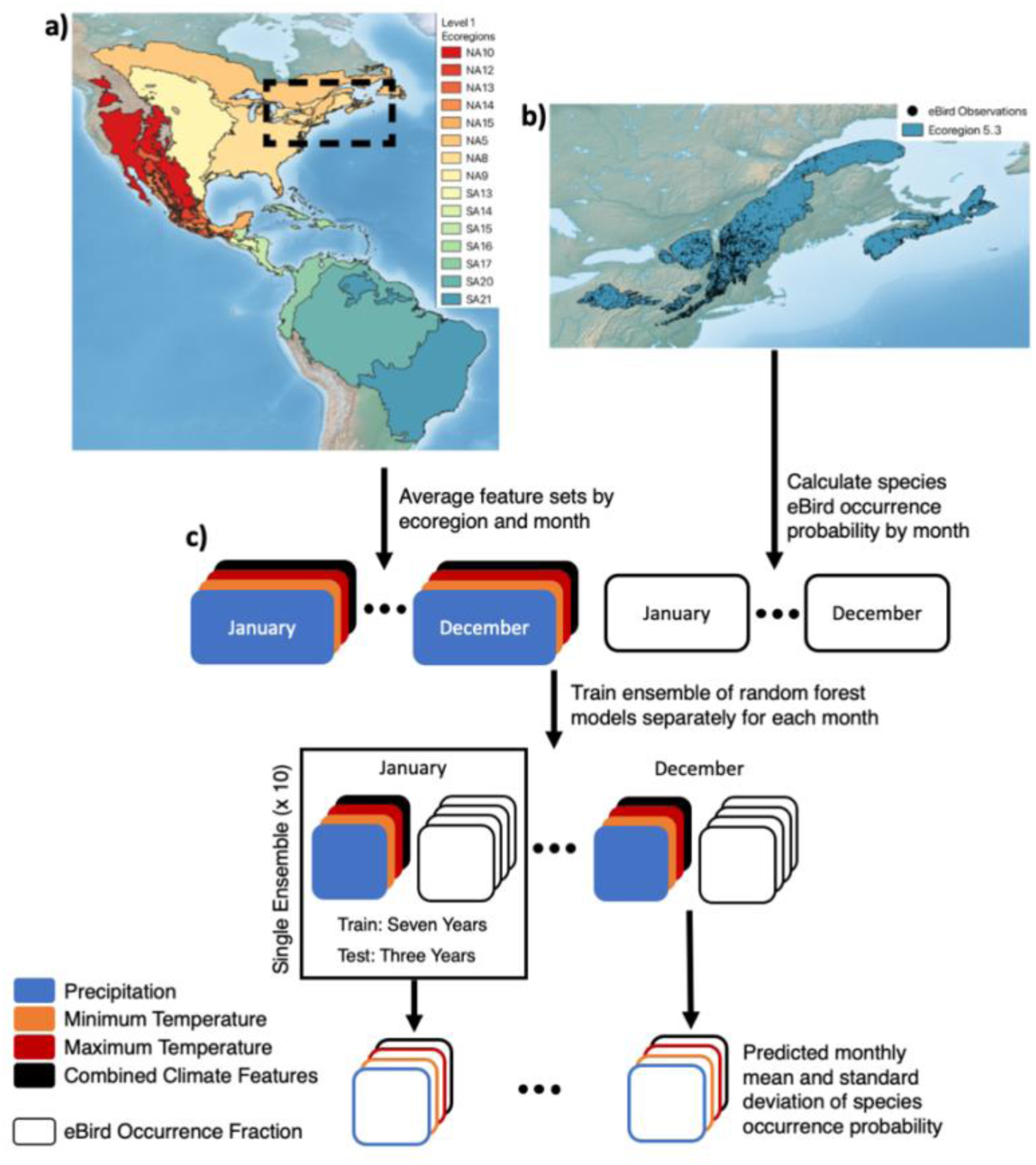
Map of targeted EPA Ecoregions and eBird survey locations. a) North and South American Level 1 Ecoregions for averaging of climate variables. b) The eBird study region Ecoregion 5.3, a subset of Level 1 Ecoregion 5 (NA5), outlined in light green with individual eBird survey coordinates in black. c) Data flow through random forest model ensembles. Monthly climate features are aggregated to the Level 1 Ecoregion spatial scale and monthly eBird species occurrence fractions across checklists are calculated for the Ecoregion 5.3 study region. An ensemble of ten models is trained and tested for each month and climate feature combination to predict checklist occurrence fractions, and the results of the ensemble are then averaged. Basemap made with Natural Earth.

### eBird Data

We used the eBird Basic Dataset from 2008-2018 ^30^ to derive the monthly occurrence rate of bird species across surveys in our study area. The dataset consists of checklists of observed species from citizen scientists, representing an “encounter rate” metric for each target species, which can be thought of as the product of that species’ detectability and occurrence probability^33^. Like other citizen science datasets, eBird checklist data can be noisy and often requires preprocessing and filtering for best results^34^. Additionally, checklists are not equally distributed spatiotemporally^35,36^. Still, eBird has been shown to closely replicate professional surveys^37^ and other modeling efforts have successfully used it to generate species distribution maps based on climate data^14^. We implement several data pre-processing steps to limit spatiotemporal variability in species detectability to create an “apparent distribution” ^33^ of encounter rate that resembles actual occurrence probability as much as possible: we restrict surveys to forested areas, only consider the most common migrating passerine species, and drop high-effort surveys. To account for increases in eBird usage throughout the study period, we also subsample surveys ^38^ annually.

Checklists were first filtered to our study Ecoregion. This spatial aggregation allowed us to limit the number of climate predictor variables in our random forest models and thus more easily study the temporal effects of regional climate on migration. Checklists were further processed according to methods recommended by Johnston et al. shown to improve the performance of species occupancy and detection models when using eBird data, especially in models with large survey sample sizes ^33^. Specifically, we included only ‘complete’ checklists, removing any surveys where an unknown species was observed, and thus retaining information on species absence. Surveys with very high effort, specifically those lasting over three hours and traveling more than 5 km, as well as area counts, were excluded to retain spatial information of observations. Group surveys, which contain redundant checklists, were condensed to a single checklist.

To provide the largest possible sampling size of observations and improve model accuracy^37^, we next limited the species of interest to the 50 most commonly occurring passerine species within the ecoregion. Passerine species generally have a lower body mass and higher cold sensitivity, which can induce migratory behaviors ^39^. Of these species, we filtered down to 41 species (mean observations: 3,571, std: 2,044) (S1 Table) which are either short-distance or long-distance migrants rather than resident species, based on distribution maps from Birds of the World ^40^ and expert opinion. To further refine our analysis to only forested regions, we filtered out all checklists with less than 50% surrounding tree cover based on 100m resolution tree cover density data ^41^. From 2008 until 2018, the number of annual surveys in eBird has greatly increased. To avoid bias in inferred species richness in later years which have a greater number of surveys, we randomly subsampled 1601 surveys at the annual scale, the number of surveys in 2008 after filtering for forest cover and survey type (S1 Fig). We then calculated the percentage of surveys on which each species appears on a monthly scale to create our target features. The final 41 target features are the mean checklist occurrence rates of the migrating passerine species of interest within forested areas of the Atlantic Highlands ecoregion.

### Climate Data

To study historical patterns in migration and climate, we utilized WorldClim monthly historical raster data ^42^ downscaled with WorldClim 2.1 ^43^ for monthly minimum temperature, maximum temperature, and precipitation at 2.5 minute spatial resolution across North and South America between 2008-2019. The raster data were averaged into EPA Level 1 Ecoregions for North and South America (Fig 1A) that fall within typical migration pathways for birds that spend their breeding season in the eBird study region. Features were then individually normalized by setting a mean of 0 and scaling features to unit variance. Each of the three feature sets (maximum temperature, minimum temperature, and precipitation) had a total of 15 features per month, from eight North American Ecoregions and seven South American Ecoregions. One feature set containing all three climate features was also created, with 45 features in total.

Similarly, to generate projected climate features, we utilized WorldClim monthly projected data from the Coupled Model Intercomparison Project Phase 6 (CMIP6) ^43^, aggregated and averaged to the same EPA Level 1 Ecoregions. The CMIP6 framework forms the standard for international climate assessments, such as those from the Intergovernmental Panel on Climate Change ^44^ and includes an ensemble of climate models from various research groups that rely on common historical training data. For each CMIP6 model, we also tested several radiative forcing pathways, dubbed the Shared Socioeconomic Pathways (SSP) ^45^, which compared the effects of various greenhouse gas emissions scenarios. Due to our use of a random forest model, which averages across previously seen training samples to make predictions and cannot extrapolate, our projections of species occurrence fall within the bounds of observed species occurrences on eBird checklists from 2008-2018. Still, to avoid projecting species occurrence rates in radically different climatic conditions, we limited our study to the 2021-2040 averaged WorldClim datasets. This consists of the mean projected monthly minimum temperature, maximum temperature, and precipitation from 2021 to 2040. We used four CMIP6 models available from WorldClim which performed best on tests comparing probability distributions of projections to historical observations and the regularity of climate fluctuations ^46^.

### Analysis of Historical Avian Migration

We trained random forests on the historical climate features to predict the eBird checklist target features (Fig 1C). Twelve ensembles of models were trained in total, one for prediction of eBird checklist occurrences in each month of the year. This separate training of models for each month of the year and lag on the climate feature dataset enabled us to study seasonal patterns in the error rates and feature importance. Each ensemble of models consists of ten 1000-tree random forest models trained with varying 70:30 train-test splits across the ten years of data being studied (2009-2018). For example, the first model was trained on 2009, 2010, and 2011 data, and tested on 2012-2018 data, the second model was trained on 2010, 2011, and 2012 data, and tested on 2013-2018 and 2009 data, and so forth. Each model is trained on 7 years x 15 aggregated climate features, or 7 x 45 when using all temperature and precipitation features. This process was then repeated while lagging the climate features from one to six months. For example, the climate features from March were used to train models to predict the checklist occurrences in April, and so forth.

From each ensemble of ten random forest models, we recorded the mean and variance of the test set predictions and mean absolute errors (MAEs) for all species. We also extracted feature importances for each climate feature and Ecoregion using the built-in random forest impurity-based metric. In regression settings, this importance reflects the total reduction in mean squared error that each feature contributes when splitting the training set across all decision trees nodes^26^. This allows us to identify which climate features are most effective for partitioning all species occurrence rates into more homogenous subsets. These importance metrics are relative, and sum to one across all features^47^. We note that impurity-based importance risks overvaluing high-cardinality features^48^. To mitigate this, we compare aggregated distributions of importances between North and South American features, rather than interpreting the importances of individual features. While permutation importance is another commonly used, model-agnostic approach, it can be heavily biased by multicollinearity between features^49^, which is prevalent in our climate feature set.

As these ensembles of models generate distributions of error and feature importance, we were able to compare their performances in permutation tests, which evaluate whether one distribution is significantly greater or less than another, allowing us to test whether certain features facilitate better model performance based on their errors, or if certain groups of features have higher importance. Specifically, we used the scipy permutation_test function^50^ with 10,000 resamples. First, we aggregated MAE across three model ensembles (max temperature, minimum temperature, precipitation) trained on climate data lagged from 0-3 months. To determine if lagged data gives lower MAE, the distribution of the lagged errors were compared to the one, two, and three month lagged distributions using permutation_test with alternative ”less”. To determine whether feature importances were higher in North American climate features than South American, we aggregated feature importances across the same three model ensembles and lags and performed permutation_test with alternative “greater” (see Table S2 for more details). Lastly, we apply Bonferroni multiple hypothesis correction to correct for multiple tested scenarios, multiplying MAE p-values by 3 and feature importance p-values by 4 to account for the number of tests.

### Comparison of Random Forest to Other Off-the-shelf Models

We compare the performance of MRF against three other typical off-the-shelf models in the sci-kit learn library ^51^: gradient boosted trees regression, support vector regression, and multiple linear regression. While the random forest model was both multivariate and multi-response, the other three architectures were limited to single response. For these architectures, separate models were trained to predict each target bird species. The tests were performed using a combined dataset of climate features (maximum temperature, minimum temperature, and precipitation) and followed the ensemble methodology outlined in the previous section. Default model parameters were used for each architecture, except for the gradient boosted trees regression, where the default number of trees was increased from 100 to 1000 to improve performance and match the number of trees used for the random forest.

### Projections of Avian Migration Shifts

We trained MRF (each with 1,000 trees)^51^ on the combined data set of 45 climate features aggregated to the Ecoregion scale (minimum temperature, maximum temperature, and precipitation) to predict the target eBird data from 2009-2015–specifically, the mean occurrence rate of 41 common migrating passerine species within forested areas of our ecoregion. Testing was performed on 2016-2018 data (S4 Fig). We then used this model to project on the 2021-2040 monthly average climate projections from the four CMIP6 models (EC-Earth3-Veg, INM-CM4-8, MPI-ESM1-2-HR, and GISS-E2-1-H.) and four SSP projections (SSP 1, 2, 3, and 5). As with the historical ensemble of models, a separate model was trained to predict each month of the year. Projected species-wise shifts in monthly occurrence probability were calculated by subtracting the mean monthly eBird checklist occurrences (2009-2018) from the mean projected occurrences for each of the four CMIP6 models and four projections.

## Results

We found accurate per-species predictions of eBird checklists for the 41 passerines studied (Fig 2A) were made with random forest using monthly mean climate data (Fig 2D) as well as with lagged climate data up to 6 months (Fig 3 E,F). Average monthly errors in predictions were generally low across all months and across climate features, ranging from 1.7 to 3.8 percent error in eBird checklist occurrence fraction averaged across all species (Fig 3, S2 Fig), with higher errors in the Spring and Summer months when species richness was also higher. Mean absolute error was lower in the winter during periods of lower occurrence for migrant species in northern areas (Fig 2 A,B). We noted a trend in the mean annual error across species where the ensembles tended to under-predict species occurrence fraction in earlier years and over-predict species occurrence fraction in later years (Fig 3A-D). This pattern in error followed a trend in the underlyingb eBird data; on average, each species in the study occurred on 2.6% fewer eBird checklists from 2009 to 2018 (Fig 2C), a trend apparent for most species (Fig 2D). We compare our proposed approach against other standard methods such as multiple linear regression, support vector machines and gradient boosted regression Multiple error metrics (mean absolute error, root mean squared error, and coefficient of determination) demonstrate that MRF performs better than these standard methods for this prediction task (S7 Fig).

**Figure 2:**
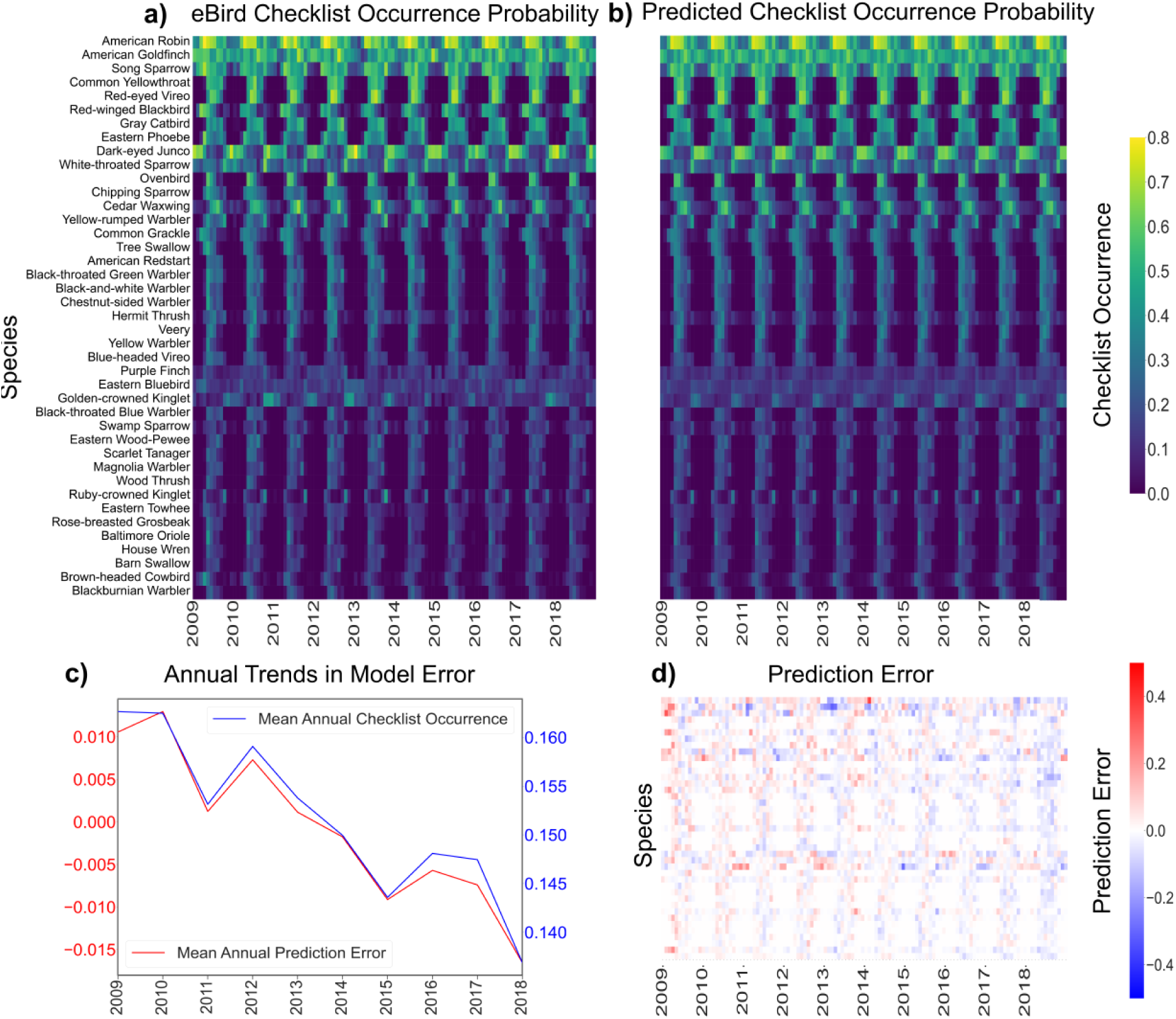
Aggregated climate data supports accurate predictions of eBird species checklist occurrences. -a) Actual eBird checklist occurrence fractions for all species. b) The random forest-predicted eBird checklist occurrence fractions for all species utilizing all available climate variables (minimum temperature, maximum temperature, and precipitation) c) Mean annual eBird checklist occurrence across all species (blue) and the mean annual error in occurrence fraction predictions (red) across all species., d) eBird checklists minus the predicted checklists using current climate feature sets, showing over-predictions in blue and under-predictions in red.

**Figure 3:**
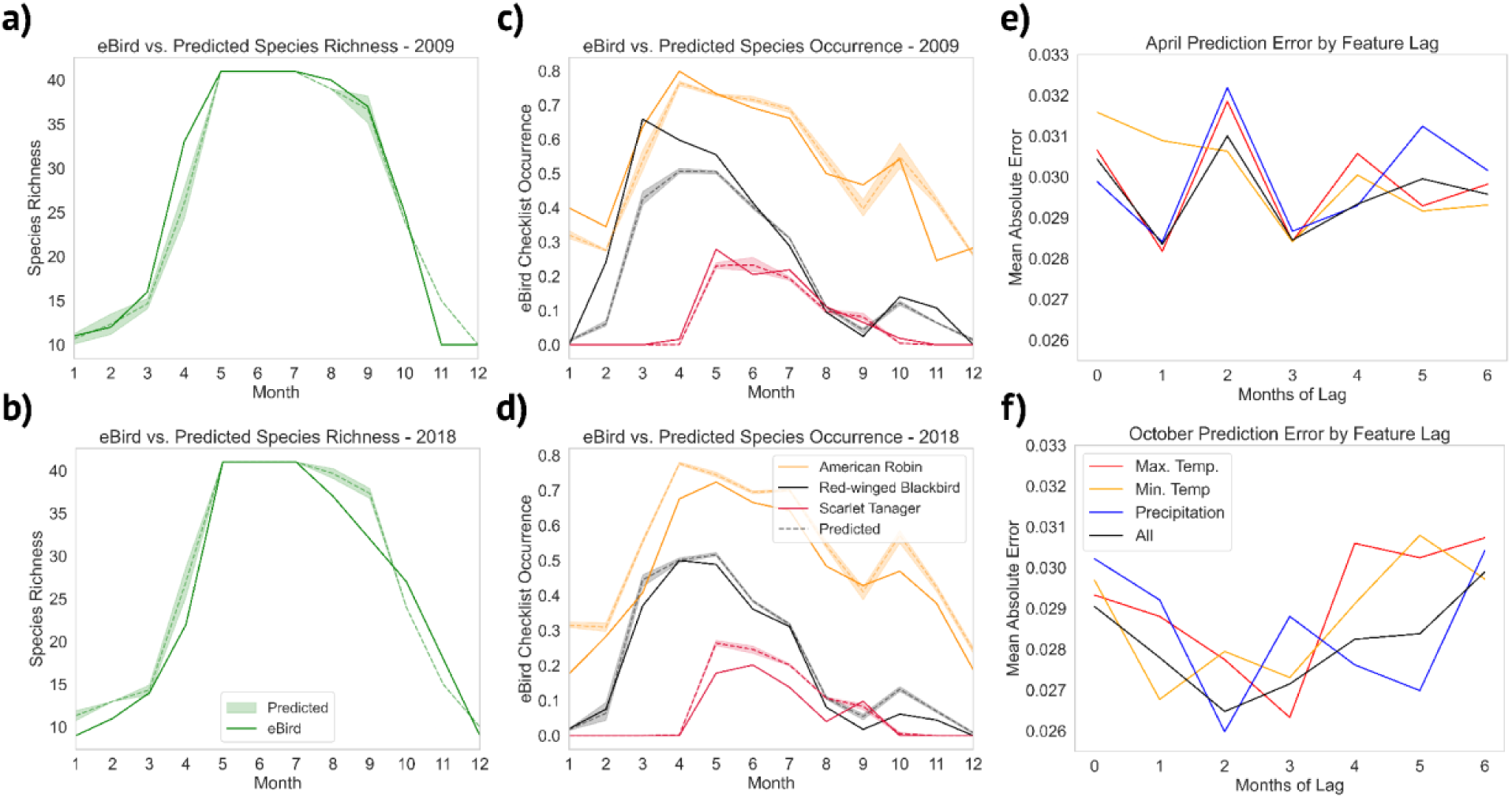
Model predictions across years suggest underlying shifts in eBird checklist occurrence and strength of lagged climate data for predictions. a) 2009 and b) 2018 eBird species richness versus mean and standard deviation of predicted species richness across the model ensemble. Species richness shown includes all species appearing on more than two percent of actual and predicted eBird checklists. c) 2009 and d) 2018 eBird ground truth versus predicted checklist occurrence fraction, showing mean and standard deviation of predictions across the model ensemble. e) Average random forest mean absolute test set errors by month of feature lag for April and f) October across all species and climate features.

Figure 3 illustrates the accuracy of the random forest ensembles for prediction of species checklist occurrence rates, but also how the temporal trends in the underlying eBird features impacted model performance. The model increasingly tended to overpredict individual species checklist occurrence towards later years in the analysis (Fig 3 C, D). Following this, the derived species richness was also increasingly over-predicted (Fig 3 A, B). Figure 3E and 3F focus on April and October specifically, as these months are transitory periods between breeding and winter seasons (Fig 2A) when migratory species are arriving and departing our study area. We noted higher volatility and no apparent trend in model error for April (Fig 3E). In October, however, the lowest error occurred when climate feature sets were lagged by one to three months (Fig 3F).

We performed permutation tests on the distributions of errors across the model ensembles to determine whether the lower error of these lagged datasets was significant. We found that lagging climate features for two and three months produced significant reductions in model MAE for predicting eBird checklist occurrences in October (p=0.0012, p=0.0024) (S2 Table). No such significant changes in error were found for lagged datasets in April. We applied permutation tests to distributions of random forest feature importance within the North and South American ecoregions (S6 Fig) to determine if the model was identifying relevant regional features. For predicting checklist occurrences in October in our northeastern study region, we found that North American climate features were significantly more important only at a lag of 3 before multiple hypothesis correction (p = 0.0221), but this result was insignificant afterwards (p=0.0884) (S2 Table). No significant spatial differences in feature importance were in predicting April occurrences.

By training on the 2008-2018 eBird checklists with all climate features and testing on CMIP6 climate projections (making broad assumptions of no change in habitat or species abundance), we projected higher positive shifts in checklist occurrence probability for April through August (ranging from 0.36% to 0.75%), with less change projected for Fall months (ranging from 0.011 to 0.19%) (Fig 4A). We found an increase in species occurrence probability across models projected for 25 of 41 species in April (mean: 0.80%), 15 species having a projected decrease (mean: -0.36%), and one species with no change (Fig 4B). Results were more mixed for October, with projected occurrence probability increases for 20 of 41 species (mean: 0.67%), decreases for 19 (mean: -0.46%), and no change for two species (Fig 4C). In contrast to the variation within eBird training data, the variance of occurrence probability for all species was effectively zero across all four CMIP6 models and four SSP projections tested, indicating that the random forest model consistently reached the bounds of known distributions of checklist occurrences across the available climate training data after the 2021-2040 period.

**Figure 4:**
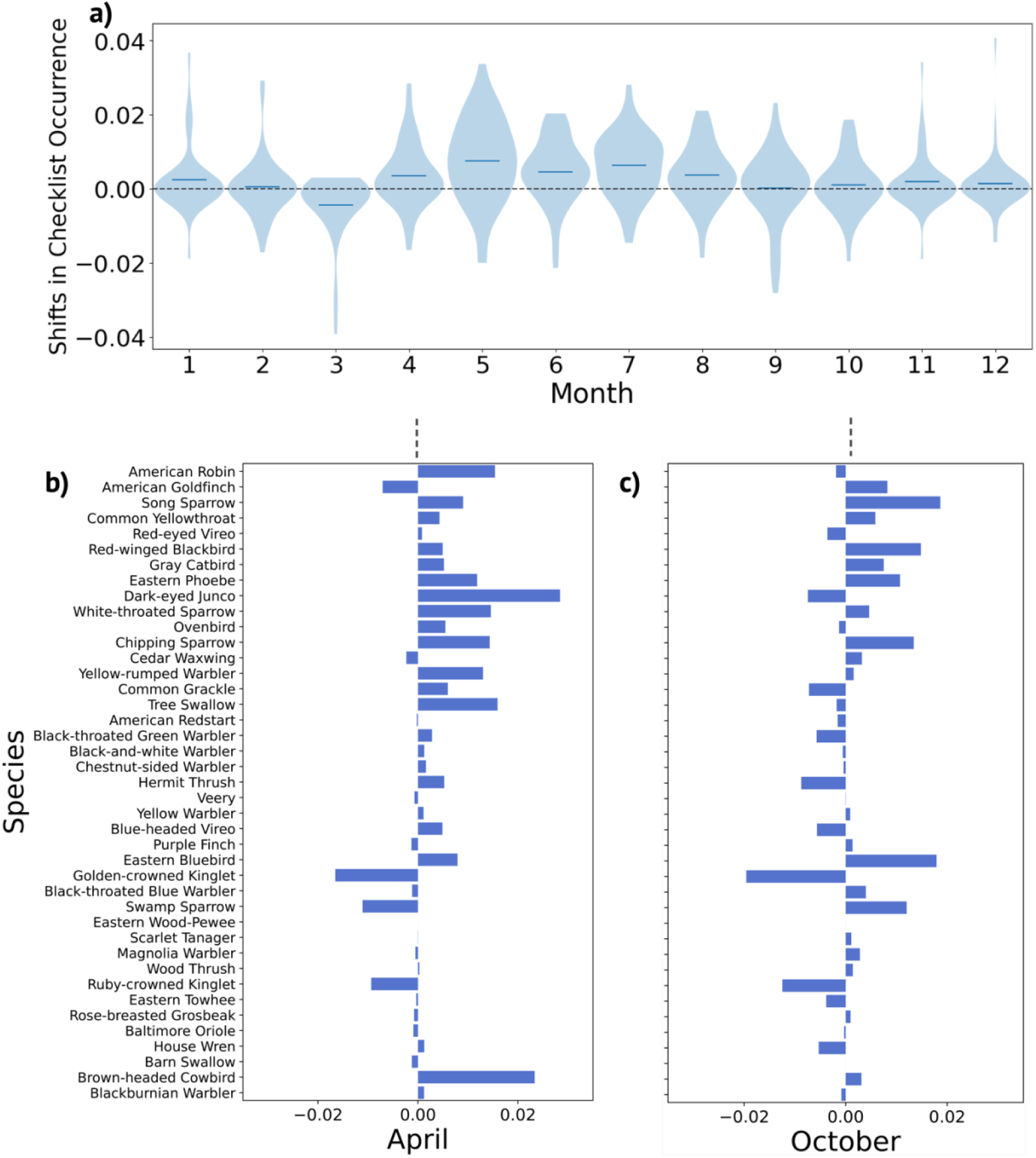
Projections of eBird checklist occurrences in a Northeastern North American ecoregion from 2021-2040 indicate higher shifts in spring occurrence probability for passerine species than in the fall. - a) Violin plot showing mean projected shifts of occurrence probability for 41 passerine species in 2021-2040 compared to historical eBird monthly means from 2008-2018 in Ecoregion 5.3 b) April and c) October per-species difference between mean historical eBird occurrence probability and mean 2021-2040 projected occurrence probability.

## Discussion

### Random Forests Accurately Predict Avian Survey Occurrence

Aggregated climate features from both North and South America allowed us to accurately predict eBird checklist occurrence probability for 41 passerine species in our North American forest study region, as well as species richness (depicted in Fig 2A and Fig 3 A, B). Importantly, the accuracy of our predictions remained consistent regardless of the specific climate variable chosen as a training feature (as illustrated in Fig 3E, 3F, and S2 Fig). Additionally, combining multiple climate features resulted in reduced variability in prediction error (as indicated in Fig 3E, F), highlighting random forest’s suitability for large, complex, feature sets in ecological settings^19^. These results are in line with previous studies which have successfully utilized eBird and climate data to generate species distribution maps using machine learning models^14,34^.

Furthermore, we observed that the trend in mean annual model error closely mirrored the trend in mean annual checklist occurrence over time across all species (as shown in Fig 2C). This alignment might be attributed to underlying patterns of decreasing biodiversity and total bird counts observed during our study period, a phenomenon that has also been documented in previous studies ^52,53^, but more study is needed to determine whether this is echoed in eBird surveys (though see Horns et al (2018)^37^). It’s worth noting that potential errors could also arise from systematic shifts in the accuracy of the underlying eBird data. However, we attempted to mitigate this risk by using randomly selected complete checklists, as recommended by Johnston et al. (2021).

### Model Metrics Capture Spatiotemporal Effects of Climate on Phenology

During October, a period when species are actively migrating southward from our study region (as depicted in Fig 3A), we observed a notable reduction in prediction error when climate features were lagged by two or three months, in comparison to using current climate features (as illustrated in Fig 3B, S2 Table). The results suggest that our model ensembles may be effectively capturing delayed temporal patterns in climate that influence migratory departure.

Furthermore, in October, the random forest analysis highlighted that North American climate features may be more influential than South American features at a lag of three months, indicating that the model may be discerning spatial climate cues contributing to migratory behavior, but this result was insignificant after applying multiple hypothesis correction.

These results echo the effectiveness of random forests in other ecological settings. In Wijeyakulasuriya et al (2020), the authors found that random forests most successfully predicted next-step movement of ants based on 39 variables describing their current state^18^. Hunt et al 2022 also applied random forests to evaluate temporal relationships between ecoclimatic variables and avian species, though using feature contribution methods instead of feature importance. They find within the model space that woodland bird species richness is heavily influenced by June vegetation indices in broadleaved forests, and that larger patches of semi-natural land reduced woodland bird diversity^20^. Overall, these findings demonstrate that use of random forests and their feature importance and error metrics can elucidate both spatiotemporal triggers for animal movement and ecoclimatic dependencies of biodiversity.

### Projections of Future Avian Phenology and Model Limitations

When projecting forward in time using models, it is essential to clearly outline scope and model limitations. Our projections of bird species from 2021-2040 were not designed to model future changes in biodiversity and population abundance. Instead, they serve the purpose of exploring how bird species might respond if sudden shifts in climatic conditions were to occur in the near future, such as within the next year. Essentially, we aimed to uncover insights from historical climate and bird migration patterns that could shed light on potential future trends. Furthermore, it’s important to note that our approach utilizes a random forest model architecture. As a result, our projections of future species occurrence probability remain constrained within the bounds of the historical training data, rendering them inherently conservative. Empirical, data-driven models such as machine learning models rely on historical observations. Thus, while these may identify ecological relationships or trends, they cannot make accurate predictions outside of historical climate covariates. These projections do not account for factors like changes in land-use and development ^10^, alterations in the timing of resource availability related to insects and flowering plants ^15,54–57^, or shifts in inter-species competition ^58^, all of which are known to exert significant influences on bird populations and distributions. Despite these limitations, we were still able to discern trends consistent with existing literature and empirical research. Overall, the model results suggest more pronounced and extensive shifts in projected future species checklist occurrences in April and May than in Fall months (as displayed in Fig 4 A-C). Indeed, many studies on shifting phenology of bird migration identify earlier Spring arrival times and breeding times with increasing temperatures ^7,15,59^, but results are less conclusive regarding shifts in autumn migration^12,17^.

### Future Work

Climate change is warming the planet unequally across geographies, disturbing evolved phenological patterns of breeding and migration ^60,61^ and contributing to population declines by inducing mismatches between arrival time and peak food availability ^15,57,59,61^. This sensitivity is especially notable in Nearctic regions ^61^, where our study area is located, and for long-distance migrants, whose wintering grounds warm at a slower rate than breeding areas ^61^. Our machine learning approach identifies similar patterns of Spring variability and sensitivity to climate change as these studies ^60,61^, which incorporate mixed linear models, while also demonstrating high predictive power for the actual occurrence probabilities of these species. The monthly scale of data resolution and the resulting small dataset size in this analysis constrained our ability to employ more deep learning techniques, such as recurrent neural networks, which potentially could capture nuanced temporal influences of climate on migration. Nonetheless, the utilization of random forests decision trees offered several advantages, including built-in feature importance methods and the ability to confine future predictions within the bounds of the training dataset. This contrasts with simpler models such as multiple linear regression, which extrapolate outside the historical range. Indeed, multiple linear regression performed poorly compared to the three tested machine learning algorithms (S7 Fig) on predicting historical data, suggesting that machine learning can improve on linear ecological models by capturing non-linear relationships with covariates ^62^.

For future research, there are several avenues to explore for enhancing our understanding of bird migration in relation to climate. One promising direction involves improving the temporal and spatial resolution of the climate datasets, which could enable machine learning models to provide a more comprehensive view of climate influence on migration patterns. Additionally, focusing on factors related to population abundance might yield valuable insights, particularly considering observed decreases in bird populations. Furthermore, incorporating other climate features that are known to affect migrations such as wind speed and direction (as highlighted by Haest et al (2019)^17^) could contribute to a more comprehensive model. Finally, the inclusion of indices derived from satellite data such as normalized difference vegetation and water indices,

which are associated with food availability and drought conditions (as mentioned by ^20,63^), could provide a more holistic understanding of the ecological factors impacting bird migrations.

## Conclusions

In this study, we present a modeling approach to identify spatiotemporal climate features that influence bird migratory behavior. Our results demonstrate that MRF models trained on these features accurately predicted eBird checklist occurrence probability throughout the year in a northern forested region. Our findings indicate that lagged feature sets were more effective in predicting passerine species occurrence probability in October compared to current climate conditions. Though statistically insignificant after accounting for multiple hypothesis correction, results suggest that models may also be assigning greater spatiotemporal importance to North American features over South American features in October, coinciding with the southward migration of species to their wintering sites. This highlights the potential of machine learning models to identify relevant spatiotemporal climate information when predicting species occurrence.

Furthermore, we explored potential future trends in migratory behavior using mean climate projections from CMIP6 for the period 2021-2040. Our results suggest an increased occurrence probability in Spring and Summer months, but a reduced effect in the Fall and Winter. These findings align with existing studies, supporting the hypothesis of a shifting of migratory timings with climate change, characterized by earlier spring arrivals and later autumn departures. Our pilot tests demonstrate that thorough investigation into model performance and feature importance within machine learning models can contribute to our understanding of complex ecological processes.

## Acknowledgements

We acknowledge the World Climate Research Programme, which, through its Working Group on Coupled Modelling, coordinated and promoted CMIP6. We thank the climate modeling groups for producing and making available their model output, the Earth System Grid Federation (ESGF) for archiving the data and providing access, and the multiple funding agencies who support CMIP6 and ESGF. SS was partially funded by a Herchel Smith Postdoctoral Research Fellowship. IAB was funded by doctoral research grant 323294 from the Research Council of Norway.

Data preprocessing computations were performed on resources provided by Sigma2–the National Infrastructure for High Performance Computing and Data Storage in Norway.

## Data Availability Statement

All data and Python notebooks required to run these analyses are available online (https://zenodo.org/records/10157596).

## Author Contributions

IAB led manuscript writing and methodology design and ran the analyses. VB, MP, KR, and SS advised on methodology design and research questions and made significant contributions to the manuscript. All authors reviewed and approved the manuscript.

