## Supplemental Information for "Exploring the Spatiotemporal Influence of Climate on American Avian Migration with Random Forests"

### Supplementary Information

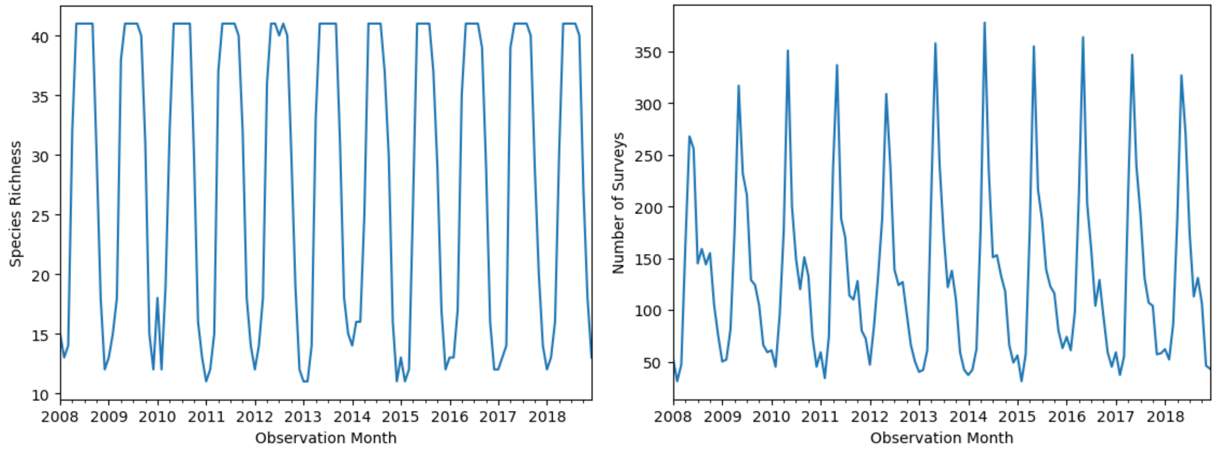


**S1 Fig. Monthly Species Richness and Survey Counts.** Monthly species richness for migrating passerine species (left) and number of sampled eBird surveys per month (right).

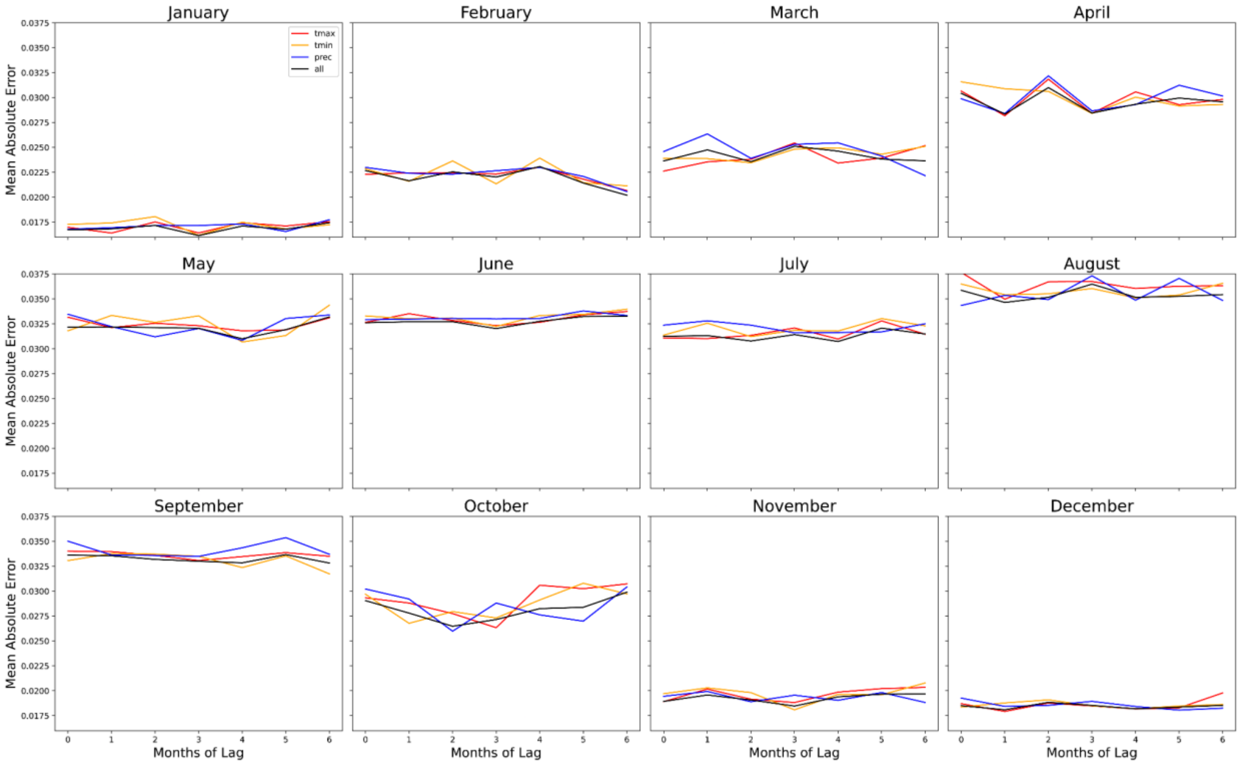

**S2 Fig. Monthly Climate Projection Model Error.** Monthly mean absolute error of eBird checklist occurrence predictions by month of feature lag and climate feature type.

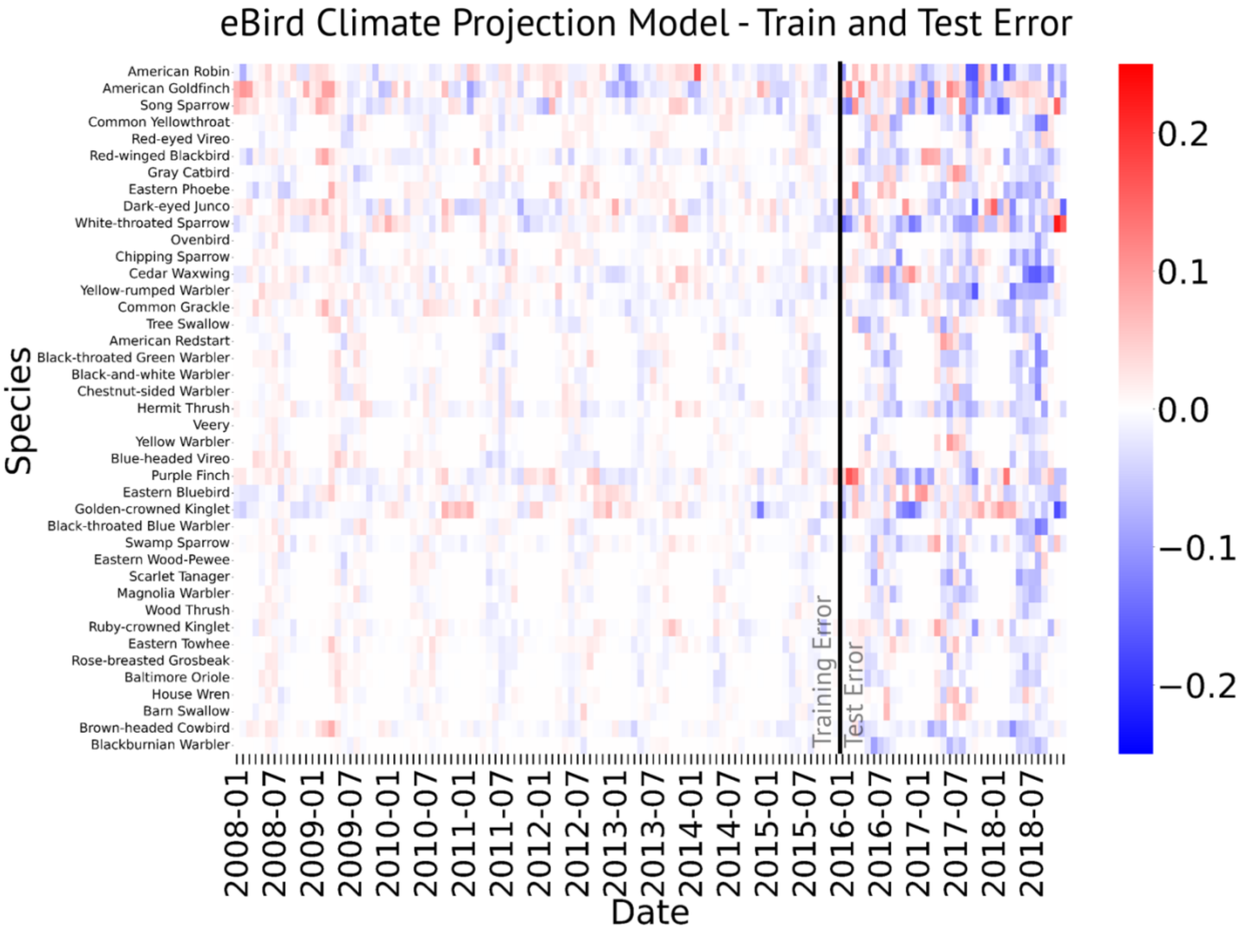


**S3 Fig. Per-Species Climate Model Error.** Random Forest training set (2008-2015) and test set (2016-2018) error for the climate projection model.

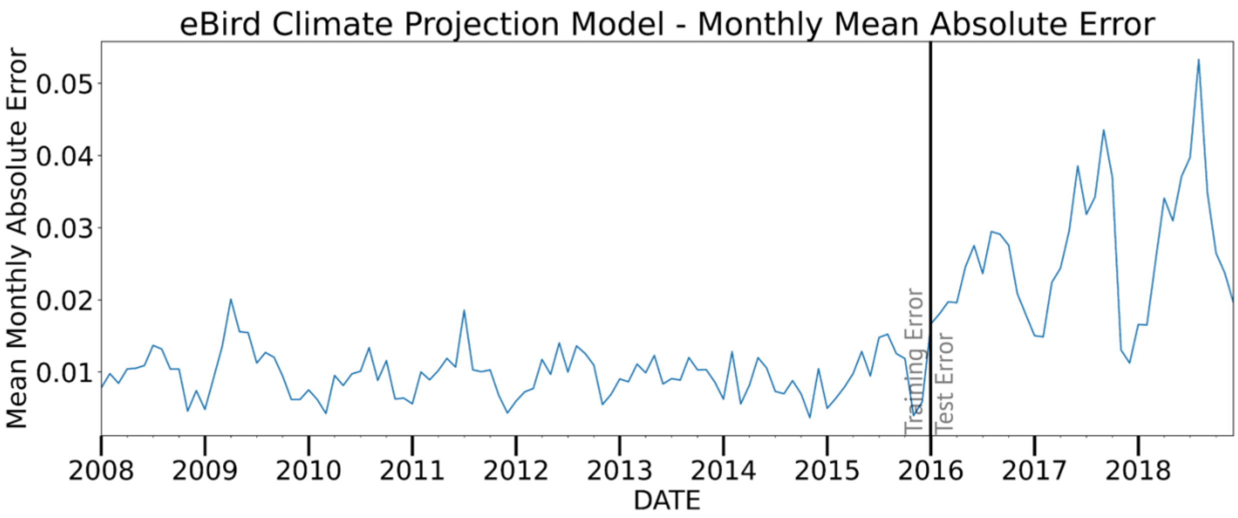


**S4 Fig. Mean Climate Model Error.** Random forest training set (2008-2015) and test set (2016-2018) monthly mean absolute error across all species for climate projection model

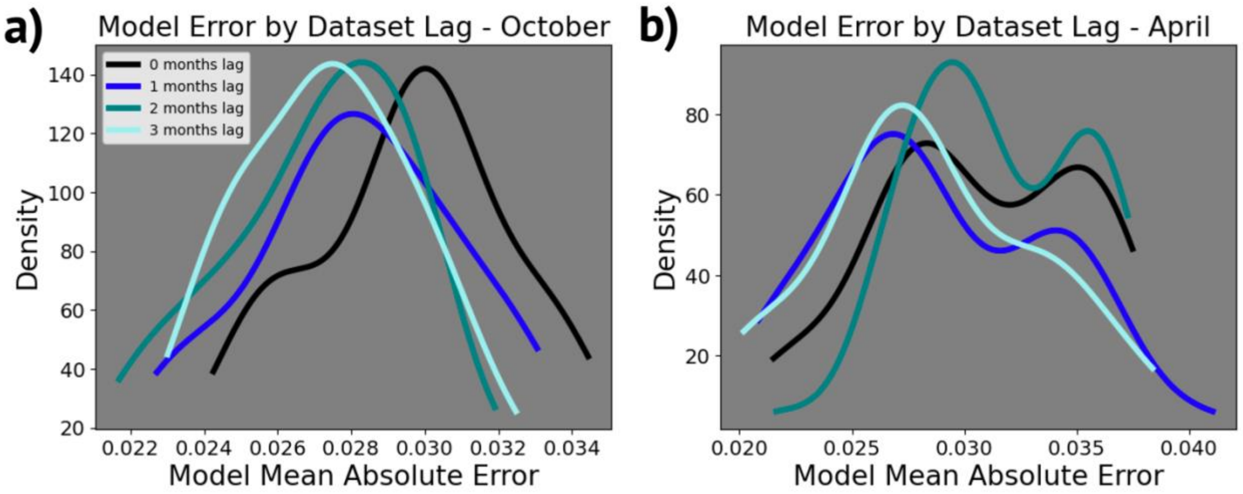


**S5 Fig. Model Error Density by Lag.** Kernel density estimation of model errors versus lag of climate features for a) October predictions and b) April predictions.

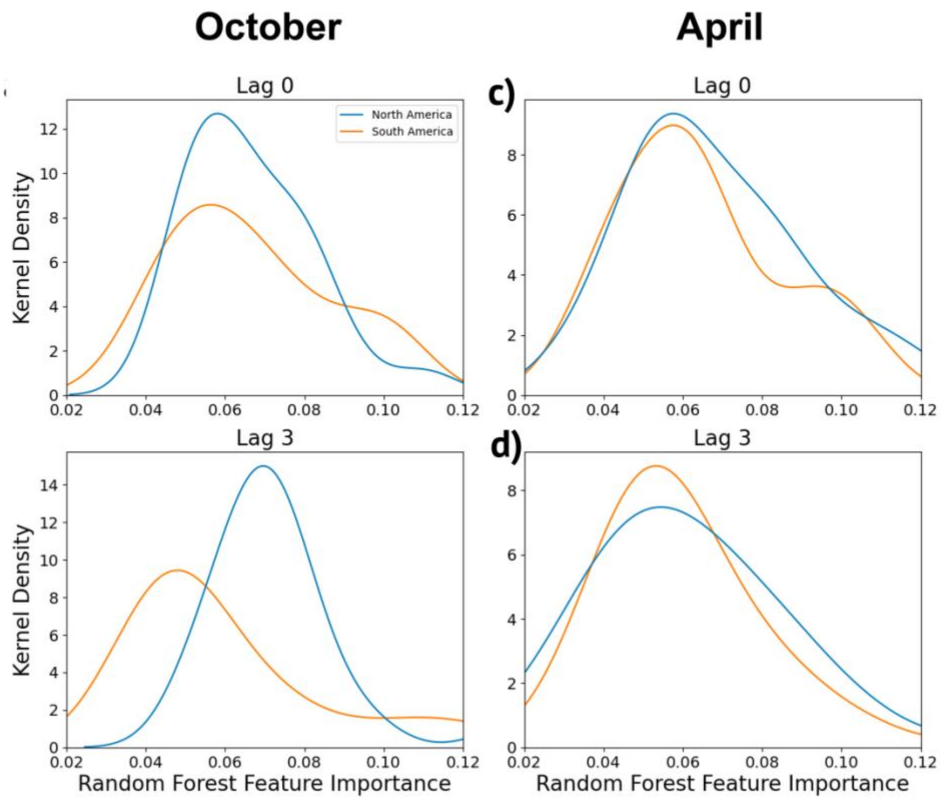


**S6 Fig. Model Feature Importance by Dataset Lag.** Kernel Density Estimation of random forest feature importance for predicting eBird checklist occurrence in a,b) October and c,d) April. Distributions are split into North (n=45) and South American (n=24) ecoregions and include model importance for each climate variable (max temperature, minimum temperature, and precipitation).

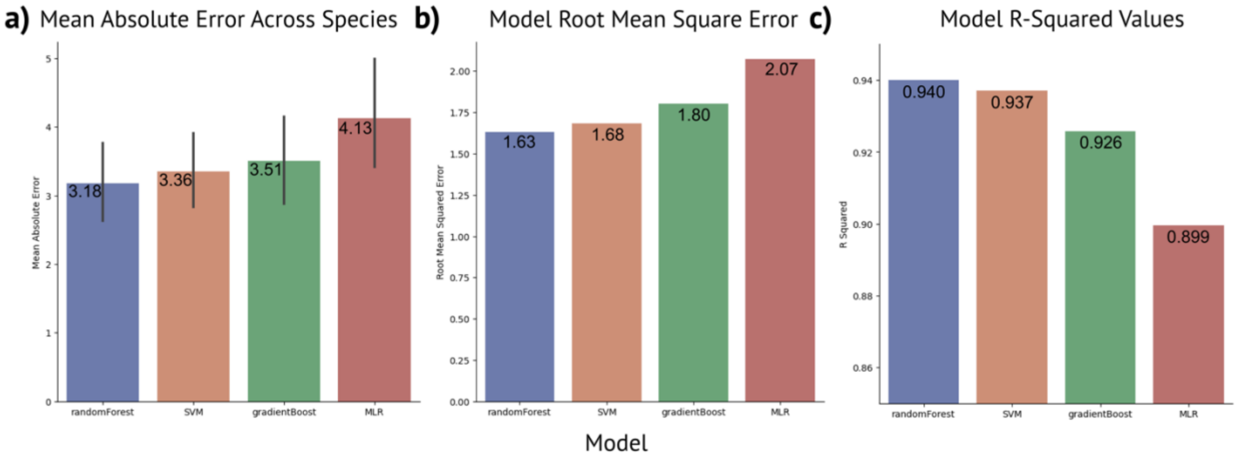

**S7 Fig. Comparison of Model Error Metrics.** a) mean absolute error, b) root mean square error, and c) r-squared across 41 predicted species between four tested models: random forest regression, multiple linear regression (MLR), gradient boosted regression trees, and support vector regression (SVM). Errors shown represent species predicted using all climate features (max temperature, minimum temperature, and precipitation) with no lag.

**S1 Table. Observation count per species across eBird checklists.**

| Species | Observations |
|---|---|
| American Robin | 10685 |
| American Goldfinch | 8637 |
| Song Sparrow | 8510 |
| Common Yellowthroat | 5881 |
| Red-eyed Vireo | 5585 |
| Red-winged Blackbird | 5491 |
| Eastern Phoebe | 5214 |
| Gray Catbird | 5184 |
| Dark-eyed Junco | 4914 |
| White-throated Sparrow | 4602 |
| Chipping Sparrow | 4464 |
| Ovenbird | 4409 |
| Cedar Waxwing | 4405 |
| Yellow-rumped Warbler | 4004 |
| Common Grackle | 3762 |
| Black-throated Green Warbler | 3371 |
| Tree Swallow | 3295 |
| American Redstart | 3177 |
| Chestnut-sided Warbler | 3011 |
| Black-and-white Warbler | 2954 |
| Hermit Thrush | 2933 |
| Veery | 2827 |
| Blue-headed Vireo | 2512 |
| Yellow Warbler | 2511 |
| Eastern Bluebird | 2418 |
| Purple Finch | 2303 |
| Black-throated Blue Warbler | 2229 |
| Scarlet Tanager | 2200 |
| Eastern Wood-Pewee | 2144 |
| Golden-crowned Kinglet | 2099 |
| Swamp Sparrow | 2067 |
| Brown-headed Cowbird | 1988 |
| Baltimore Oriole | 1987 |
| Rose-breasted Grosbeak | 1974 |
| Eastern Towhee | 1933 |
| Wood Thrush | 1911 |
| Magnolia Warbler | 1900 |
| House Wren | 1837 |
| Blackburnian Warbler | 1787 |
| Barn Swallow | 1764 |
| Ruby-crowned Kinglet | 1689 |

**S2 Table. Permutation Tests.** Permutation test results comparing distributions of model mean absolute errors for eBird checklist occurrences in a) October and b) April. Model errors across several lags of the climate datasets (max temperature, minimum temperature, precipitation) are compared to model errors with no lag in the datasets. P-values determine if the lagged results are in a distribution that is significantly less than the no lag error distribution (see Figure S7). A similar process is applied on the distributions of random forest feature importance (see Figure S8) to determine if North American climate features (n=45) are significantly more important than South American climate features (n=24) for predicting eBird checklist occurrence in c) October and d) April. Multiple hypothesis corrections are applied to each p-value to correct for multiple experiments in each month; each MAE p-value is multiplied by three to account for the three lags tested, and each feature importance p-value is multiplied by four to account for the four comparisons between North and South American feature importances.

| a) October MAE – Permutation Test (n=30) | | | |
|---|---|---|---|
| Lag | Test Statistic | p-value | Corrected p-value |
| 0 | 0 | 0.5000 | --- |
| 1 | 0.0015 | **0.0227** | 0.0681 |
| 2 | 0.0025 | **0.0004** | **0.0012** |
| 3 | 0.0023 | **0.0008** | **0.0024** |

| b) April MAE – Permutation Test (n=30) | | | |
|---|---|---|---|
| Lag | Test Statistic | p-value | Corrected p-value |
| 0 | 0 | 0.4960 | --- |
| 1 | -0.0016 | 0.1140 | 0.3420 |
| 2 | 0.0009 | 0.7640 | 2.292 |
| 3 | -0.0022 | **0.0396** | 0.1188 |

| c) October Importance – Permutation Test (n=(45,24)) | | | |
|---|---|---|---|
| Lag | Test Statistic | p-value | Corrected p-value |
| 0 | -0.0002 | 0.5232 | 2.0928 |
| 1 | -0.0027 | 0.6922 | 2.7688 |
| 2 | -0.0106 | 0.9673 | 3.8692 |
| 3 | 0.0124 | **0.0221** | 0.0884 |

| d) April Importance – Permutation Test (n=(45,24)) | | | |
|---|---|---|---|
| Lag | Test Statistic | p-value | Corrected p-value |
| 0 | 0.0039 | 0.3032 | 1.2128 |
| 1 | -0.0013 | 0.5686 | 2.2744 |
| 2 | -0.0068 | 0.8283 | 3.3132 |
| 3 | 0.0056 | 0.2817 | 1.1268 |
